# Post-transcriptional modulation of the SigF regulon in *Mycobacterium smegmatis* by the PhoH2 toxin-antitoxin

**DOI:** 10.1101/2020.01.15.907535

**Authors:** Emma S.V. Andrews, Elizabeth Rzoska-Smith, Vickery L. Arcus

## Abstract

PhoH2 proteins are highly conserved across bacteria and archaea yet their biological function is poorly characterised. We examined the growth profiles of *Mycobacterium smegmatis* strains *mc*^*2*^*155* and *mc*^*2*^*155 ΔphoH2* and observed the same growth profile and growth rate in a variety of conditions. In light of the comparable growth rates, we used RNAseq to provide a snapshot of the differences between the transcriptomes of *M. smegmatis mc*^*2*^*155* and *M. smegmatis mc*^*2*^*155 ΔphoH2* during normal growth. At 48 hours, elevated expression of the *sigF* regulon and its predicted regulatory cascade was observed in *ΔphoH2* relative to wild type. In biochemical assays, PhoH2 showed specific activity toward *sigF* mRNA insinuating a role of PhoH2 in modulating the pool of *sigF* mRNA in the cell during normal growth, adding further complexity to the repertoire of reported mechanisms of post-translational regulation. Multiple copies of the preferred target site of PhoH2 were identified in loops of the *sigF* mRNA structure, leading us to propose a mechanism for the activity of PhoH2 that is initiated after assembly on specific single-stranded loops of RNA. We hypothesise that PhoH2 is a toxin-antitoxin that contributes to the regulation of SigF at a post-transcriptional level through targeted activity on *sigF* mRNA. This work presents the first evidence for post-transcriptional regulation of SigF along with the biological function of PhoH2 from *M. smegmatis*. This also has implications for the highly conserved PhoH2 toxin-antitoxin module across the mycobacteria including the important human pathogen *M. tuberculosis*.

## Introduction

Sigma factors initiate gene expression through their interaction with RNAP [1]. Their function directs the binding of RNAP to specific promoter sites to initiate transcription of specific subsets of genes [1]. In mycobacteria, sigma factors SigA and SigB are responsible for the expression of essential genes [2]. Alternate sigma factors function to coordinate gene regulation in response to different environmental stresses and changing physiological conditions [3]. One such alternate sigma factor, SigF, is involved in the cell’s adaptation to stationary phase, heat, oxidative stress and antimicrobials [4-6]. SigF has been shown to regulate cell wall composition through the modulation of lipid biosynthesis [7] suggesting a prominent role in mycobacterial cell wall structure and function, persistence and pathogenesis [8]. Evidence in the literature points towards complex, post-translational regulation of SigF, through the activity of anti-sigma factors and their antagonists [6, 7, 9] although there are no reports regarding regulation at the post-transcriptional level.

PhoH2 is a modular enzyme with a PIN-domain RNAse fused with a PhoH-domain RNA helicase [10, 11]. In mycobacteria, PhoH2 is co-transcribed with a short, upstream gene, *phoAT*, whose protein product interacts with PhoH2 [10]. This small protein is comparable to VapB of VapBC (PIN-domain) toxin-antitoxin systems, where VapB functions as a transient inhibitor of VapC (PIN-domain) [12]. Experimental investigations show that PhoH2 from *Mycobacterium tuberculosis* and *Mycobacterium smegmatis* has sequence-specific RNAse and ATP-dependent helicase activities, with preference for double-stranded RNA that contains a 5’-3’ overhang, and where the terminal combination of RNA bases is 5’-A C [A/U] [A/U] [G/C] [10]. This suggests that PhoH2, similar to other PIN-domain containing proteins [12], is a toxin-antitoxin with additional RNA helicase activity that acts on specific RNA substrates and is likely to play a role in the adaptive response to changing environmental conditions [10, 11].

Outside the mycobacteria PhoH2 has been shown to play a role in regulating the infection process of *Synechococcus sp. strain WH8102* by Cyanomyovirus [13]. In *Corynebacterium glutamicum*, PhoH2 is implicated in the response to phosphate limitation [14] and there is evidence for reduced expression of *phoH2* during the transition from exponential growth to stationary phase growth [15]. This suggests that the expression of *phoH2* is not under control of the prominent sigma factor, SigB (analogous to SigF from *M. smegmatis*) during this phase of growth.

To determine the biological function of PhoH2 from *M. smegmatis* we initially examined growth profiles of *M. smegmatis mc*^*2*^*155* and *M. smegmatis mc*^*2*^*155 ΔphoH2* under rich, defined and nutrient limiting conditions. Under standard rich conditions, RNAseq was performed on cells from both strains harvested at points along the growth curve. At 48 hours, at the onset of stationary phase, the greatest difference to the expression profile between *M. smegmatis mc*^*2*^*155 ΔphoH2* and *M. smegmatis mc*^*2*^*155* was observed. The majority of genes upregulated in *ΔphoH2* compared to wild type were those belonging to the SigF regulon and its predicted regulatory cascade. This suggested dis-regulation of SigF and its associated genes in the absence of PhoH2. To investigate the involvement of PhoH2 in the regulation SigF and its regulatory cascade, biologically relevant RNA transcripts were used as targets in RNAse assays with PhoH2. PhoH2 showed specific activity toward *sigF* mRNA and we identified multiple copies of the preferred target site of PhoH2 within loops of the predicted *sigF* mRNA structure. This suggested that PhoH2 assembles and initiates its activity on specific single-stranded loops of mRNA. This work presents the first evidence for post-transcriptional regulation of SigF along with the biological function of PhoH2 from *M. smegmatis* and predicted mechanism for PhoH2 activity.

## Materials and Methods

### Construction of ΔphoH2 knockout in *M. smegmatis mc*^*2*^*155*

An unmarked deletion of *phoH2* was created by a two-step allelic exchange mutagenesis [16]. For this purpose a construct containing 822 bp regions flanking the *phoH2* gene on the left and right respectively (using primers listed in Table S1), was cloned into pX33 to yield pX33-*phoH2* LFRF. This plasmid was transformed into *M. smegmatis mc*^*2*^*155* and transformants were selected at 28 °C in the presence of 5 mg/ml gentamycin. For deletion of *phoH2*, strains carrying pX33-*phoH2* LFRF were grown in the presence of gentamycin at 42 °C to select for integration of the plasmid into the chromosome of *M. smegmatis mc*^*2*^*155* via a single crossover event. Colonies were screened for integration by exposure to 250 mM catechol. Selected colonies were grown in LBT medium at 37 °C and aliquots of these cultures were plated onto low salt (2 g/l NaCl) LBT plates containing 10 % sucrose and incubated at 42 °C to select for a second crossover event leading to the loss of the plasmid and deletion of *phoH2*. Colonies were screened for loss of the plasmid with 250mM catechol and candidate mutants were screened by PCR using primers that flanked the deletion site.

### Growth of *M. smegmatis mc*^*2*^*155* and *mc*^*2*^*155 ΔphoH2* in rich, defined and nutrient limiting conditions

*M. smegmatis* strains *mc*^*2*^*155* and *mc*^*2*^*155 ΔphoH2* were grown in LB media containing a final concentration of 0.05 % tyloxapol. Three overnight starter cultures in LB media grown overnight at 37 °C 200 rpm were used to seed three cultures at a starting OD_600_ of 0.01. For defined and nutrient limiting experiments, the overnight LB starter cultures were used to seed a second defined starter culture (Modified Sautons - 0.5g/L MgSO_4_.7H_2_0, 2 g/L citric acid, 1g/L L-asparagine, 0.3 g/L KCl.H_2_0, 0.2 % glycerol, 0.64 g/L FeCl_3_, 100 µM NH_4_Cl and 0.7 g/L K_2_HPO_4_.3H_2_0). The second starters were incubated overnight at 37 °C 200 rpm. These were used to seed three cultures of defined and/or nutrient limiting media, at a starting OD_600_ of 0.01. For nutrient limiting cultures, the carbon, nitrogen or phosphate source was reduced to 0.05 %, 0.05 g/L and 40 µM respectively. Cultures were incubated for up to 120 hours (5 days) and growth was monitored by optical density (OD_600_) at regular intervals and curves plotted and analysed for significance using an unpaired t-test (p=0.05) using Prism V7.

### RNA isolation and sample preparation for RNAseq

Cells from each of the three cultures were harvested for RNA isolation at 24, 48 and 72 hours. These were immediately combined with 5 M GITC at a 1:4 ratio of cells to GITC. These were spun down and resuspended in 0.5 ml 5 M GITC and stored in a tube containing approximately 0.3 g 0.1 mm and 2.5 mm zirconia beads. Cells were ruptured using a Fast Prep cell disrupter (FP120 Thermo Savant) for increasing time periods. *The cells were incubated at RT with 50 µl 2 M sodium acetate pH 4 for 10 minutes, before incubation on ice with 100 µl 1-bromo, 3-chloro propanate for 5 minutes. Samples were spun to separate the phases and the top layer collected and the process repeated twice from *. The final top layer was incubated at −40 °C overnight with an equal volume of isopropanol. Samples were spun at 13000 rpm for 15 minutes at 4 °C. The precipitated RNA was dissolved in 100 µl 10 mM Tris-HCl pH 7, 0.5 mM MnCl_2_ and DNAse treated with 2.5 µl Promega DNAse for 30 minutes at 37 °C. The samples were incubated in a solution of 5.2 M guanidium thiocynate, 2 M guanidine HCl and 2 M urea for 5 minutes at RT, prior to incubation for 10 minutes at RT with 400 µl isopropanol. Samples were spun at 13000 rpm for 15 minutes at RT, the pellet was washed with 70 % ethanol then dissolved in 200 µl of RNAse free H_2_0. To this, 200 µl of 5 M LiCl_2_ was added and incubated for 1 hour at −20°C. Samples were spun at 13000 rpm for 15 minutes at 4 °C and the pellet resuspended in 100 µl RNAse free H_2_O. The RNA was precipitated with 10 µl 3 M sodium acetate pH 5.2 and 275 µl 100 % ethanol at −20 °C for 10 minutes. Samples were spun at 13000 rpm for 15 minutes at 4 °C. The pellet was washed with 70 % ethanol and spun at 13000 rpm for 15 minutes at 4 °C. The final RNA pellet was resuspended in 25 µl 10 mM Tris-HCl pH 7, 0.5 mM MnCl_2_. The ‘best’ RNA samples as determined by A260/A280 and A260/A230 ratios and gel analysis from each strain/time point were pooled and sent for sequencing in 75 % ethanol.

### Transcriptome analysis

The transcriptome of each RNA sample was sequenced at the Beijing Genomics Institute (BGI), China. RNA samples that met library construction requirements (RIN >0.8) had their rRNA removed and were fragmented for cDNA synthesis for sequencing on an Illumina-HiSeq2000/2500. Raw reads were filtered and the clean reads aligned with the genome of *M. smegmatis mc*^*2*^*155* (NC008596.1) using SOAPaligner/SOAP2. The alignment was used to calculate the distribution of reads and coverage. An initial table of differentially expressed genes between *M. smegmatis mc*^*2*^*155* and *M. smegmatis mc*^*2*^*155 ΔphoH2* was compiled that had a FDR ≤ 0.001 and an absolute value of Log2 ratio of ≥1. These genes were further manually curated and shortlisted based on the following criteria ≥2 Log2 ratio and ≥75 reads.

### Protein expression and purification

PhoH2 and PhoH2-R339A were expressed and purified as described in Andrews & Arcus (2015) [10]. Briefly, a single colony was used to inoculate an LB seeder culture supplemented with 50 mg/ml kanamycin. This culture was grown for 24 h at 37 °C and was used at a 1:100 dilution to inoculate an LB expression culture supplemented with 50 mg/ml kanamycin. These cultures were incubated at 37 °C and were induced with a final concentration of 0.75 mM IPTG at an OD_600_ of 0.4-0.6 and further incubated with shaking at 37 °C overnight. Cells from large-scale expression cultures were harvested. For purification, cells were resuspended in lysis buffer, 50 mM TRIS pH 8, 200 mM NaCl 5 mM MgCl_2_, sonicated on ice and harvested by centrifugation. The soluble fractions containing His-tagged PhoH2 or PhoH2-R339A were purified by IMAC on a HisTrap HP column (GE Healthcare, UK). The protein fractions were purified further by size exclusion chromatography, using an S200 10/300 Superdex™ column (GE Healthcare, UK) in the same buffer.

### Biological target assays

Target DNA sequences, *sigF, rsbW-sigF* and *chaB-rsbW-sigF* were amplified from *M. smegmatis mc*^*2*^*155* genomic DNA using SF Fwd and Rev, RS Fwd and SF Rev, and UCRS Fwd and SF Rev respectively (Table S1) in PCR reactions with either KAPA HiFi DNA polymerase with high GC buffers (KAPA Biosystems) or Hot FIREPol Blend Master mix (Solis BioDyne). These products were used as DNA template for a second round of PCR using T7+SF Fwd, T7+RS Fwd and T7+UCRS Fwd in place of the original forward primer to introduce the T7 promoter sequence to the 5’ end of the PCR product. The resulting +T7 PCR products were transcribed into RNA using MEGAscript T7 transcription kit (Thermo Fisher Scientific). Purified PhoH2 (125 µM) was incubated with target RNA (25 µM) in a reaction containing 5 mM ATP made up to 15 µl with assay buffer (50 mM tris pH 7.5 20 mM NaCl 5 mM MgCl_2_) for 5, 15 and 30 minutes at 37 °C. Reactions were quenched with and equal volume of 2 x formamide stop solution (80% formamide (v/v), 5 mM EDTA, 0.1% (w/v) bromophenol blue). Prior to analysis by electrophoresis on a 1x TAE 1.5 % gel, reactions were heated at 70 °C for 5 min and cooled immediately on ice. The results were visualised by staining with 1x SYBR safe nucleic acid stain (Invitrogen).

## Results

### *M. smegmatis mc*^*2*^*155 ΔphoH2* adopts the same growth profile as *M. smegmatis mc*^*2*^*155* in rich, defined and nutrient limiting growth conditions

To screen for differences in the growth profile of *M. smegmatis mc*^*2*^*155 ΔphoH2* (Fig S1) compared with its parent *M. smegmatis mc*^*2*^*155*, strains of *M. smegmatis* were grown in standard rich, defined and nutrient limiting conditions and growth measured by optical density (OD_600_). Fig 1 shows that *M. smegmatis mc*^*2*^*155 ΔphoH2* adopts the same growth profile as its parent in standard rich, defined and nutrient limiting conditions and that the growth rates also match (Table 1).

**Table 1:** Growth rates of *M. smegmatis mc*^*2*^*155* and *M. smegmatis mc*^*2*^*155 ΔphoH2* in rich, defined and limiting growth media. The growth rate (G) of each culture was calculated using OD_600_ measurements from two time points (B and b) in exponential phase, where G = t/3.3 logb/B. Growth rate is presented in hours as the mean of three cultures of each strain ± SD.

| Growth media | G (hours)<br>*M. smegmatis*<br>*mc<sup>2</sup>155* | G (hours)<br>*M. smegmatis*<br>*mc<sup>2</sup>155 ΔphoH2* |
| --- | --- | --- |
| LB - rich | 2.80 ± 0.07 | 2.80 ± 0.04 |
| Sautons - defined | 3.95 ± 0.14 | 3.82 ± 0.01 |
| Sautons - carbon limiting | 6.99 ± 0.44 | 7.08 ± 0.81 |
| Sautons - nitrogen limiting | 3.99 ± 0.04 | 3.84 ± 0.08 |
| Sautons - phosphate limiting | 11.61 ± 1.70 | 12.76 ± 0.92 |

**Fig 1:**
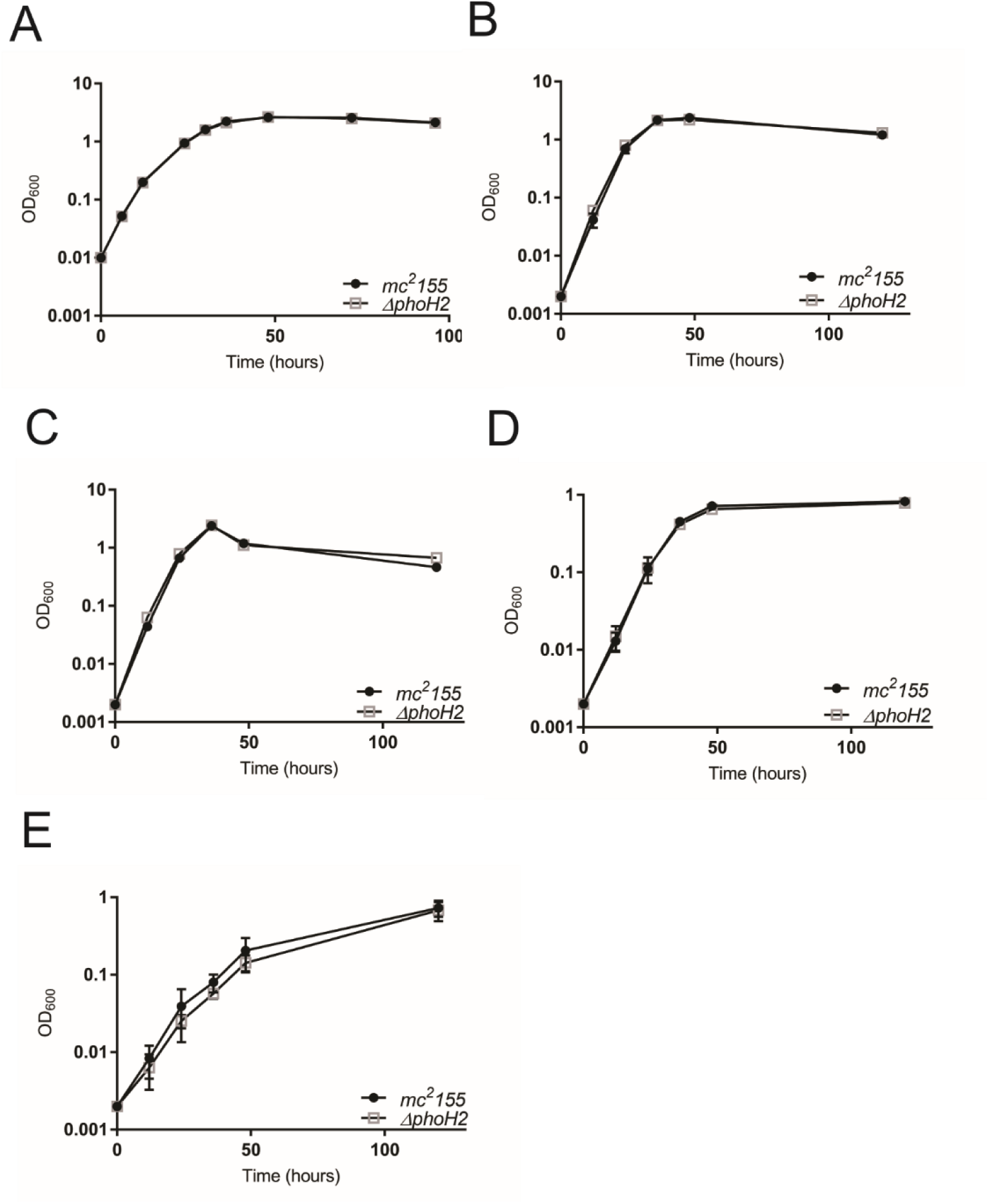
*M. smegmatis mc*^*2*^*155 ΔphoH2* adopts the same growth profile as *M. smegmatis mc*^*2*^*155* in rich, defined and nutrient limiting growth conditions. *M. smegmatis mc*^*2*^*155* (closed circles) and *M. smegmatis mc*^*2*^*155 ΔphoH2* (open squares) were cultured in (A) LB, (B) Sautons, (C) Sautons - carbon limiting, (D) Sautons -nitrogen limiting and (E) Sautons - phosphate limiting media. Growth was measured by monitoring optical density (OD_600_) at regular intervals. Data were plotted as the mean and SD of three biological replicates of each strain and an unpaired t-test was used to test for significant differences between the strains. Plots show that *M. smegmatis mc*^*2*^*155 ΔphoH2* adopts the same growth profile as its parent in rich, defined and nutrient poor conditions. (Plots generated and analysed in Prism V7).

Both strains reached higher optical densities when growth in rich, defined and carbon limiting media than when grown in limiting nitrogen or phosphate media, indicating stunted growth when these nutrients are deficient. Growth was approximately 3x slower in phosphate limiting conditions compared with when phosphate was sufficient, further suggesting growth limitations when phosphate is reduced.

### RNAseq reveals an upregulation of SigF regulon genes in *M. smegmatis mc*^*2*^*155 ΔphoH2* at 48 hours

Due to the comparable growth profiles between *M. smegmatis mc*^*2*^*155* and *M. smegmatis mc*^*2*^*155 ΔphoH2* and identical growth rate in rich media, we used RNAseq to provide a snapshot of the differences between the transcriptomes of *M. smegmatis mc*^*2*^*155* and *M. smegmatis mc*^*2*^*155 ΔphoH2.* Cells of each strain were harvested after 24, 48, and 72 hours of growth and RNA was isolated from cells harvested from each of the three cultures and the ‘best’ RNA, as determined by A260/A280 and A260/A230 ratios and gel analysis were pooled and stored in 75 % ethanol. The transcriptome sequencing and downstream analysis for differentially expressed genes (DEGs), were performed at the Beijing Genomics Institute (BGI). RNA harvested from cells upon entry into stationary phase (48 hours of growth) revealed the greatest change in gene expression between *M. smegmatis mc*^*2*^*155 ΔphoH2* and *M. smegmatis mc*^*2*^*155* (FDR ≤ 0.001 and absolute value of Log2 ratio of ≥1). These genes were further manually shortlisted (≥2 fold expression change Log2 ratio of ≥2 and ≥75 reads) and 89 genes were upregulated in *M. smegmatis mc*^*2*^*155 ΔphoH2* and 1 was downregulated compared with *M. smegmatis mc*^*2*^*155* (File S2).

Of the 89 upregulated genes, 80 belonged to the SigF regulon [6, 17] accounting for 70 % of known SigF regulated genes exclusive to stationary phase [6] and 90 % of all the upregulated genes identified in this study. The downregulated gene (MSMEG_0586), is a predicted anti-sigma factor antagonist. Of the remaining SigF regulon genes, one did not meet shortlist criteria (≥2 fold expression change, ≥75 reads) but showed an increase in expression, and the other remaining genes did not show differential gene expression in this study. The genes that were upregulated in this study, that were not identified to be part of the SigF regulon, encode for hypothetical proteins, an antigen 85-C protein and a cluster of genes (MSMEG_1974 - 1979).

### PhoH2 is involved in the regulation of the predicted SigF regulatory cascade

The genomic arrangement of *sigF* is conserved among mycobacteria [4]. In *M. tuberculosis sigF* is positioned downstream of anti-sigma factor gene, *usfX* and both genes are transcribed from a *sigF* dependent promoter positioned immediately upstream of *usfX* [4]. The genes controlled by SigF constitute the SigF regulon and these genes share a consensus promoter sequence (NGNTTG-N_14-18_-GGGTAT) [8]. In non-pathogenic *M. smegmatis, sigF* is co-transcribed with a predicted anti-sigma factor RsbW (MSMEG_1803) and a protein of unknown function, ChaB (MSMEG_1802), from two promoter sites; one upstream of MSMEG_1802, and one upstream of *rsbW*, similar to *usfX* in *M. tuberculosis* [18] (Fig 2A). Expression from the promoter positioned upstream of *chaB* is dependent on SigF and shows a 2-fold increase in transcription upon entry into stationary phase (Fig 2B) but not under conditions of stress including heat shock, acidic pH and oxidative stress [18]. Expression from the second promoter position, upstream of *rsbW*, is independent of SigF and is constitutive throughout growth and exposure to stress [18]. Expression from this position in *M. tuberculosis* is also constitutive, however unlike in *M. smegmatis*, an increased level of expression is observed upon entry into stationary phase and stress conditions [3, 5]. Like SigF from *M. tuberculosis*, the regulon of genes controlled by SigF in *M. smegmatis* share a consensus promoter sequence (GTTT-N_(15-17)_-GGGTA) [6].

**Fig 2:**
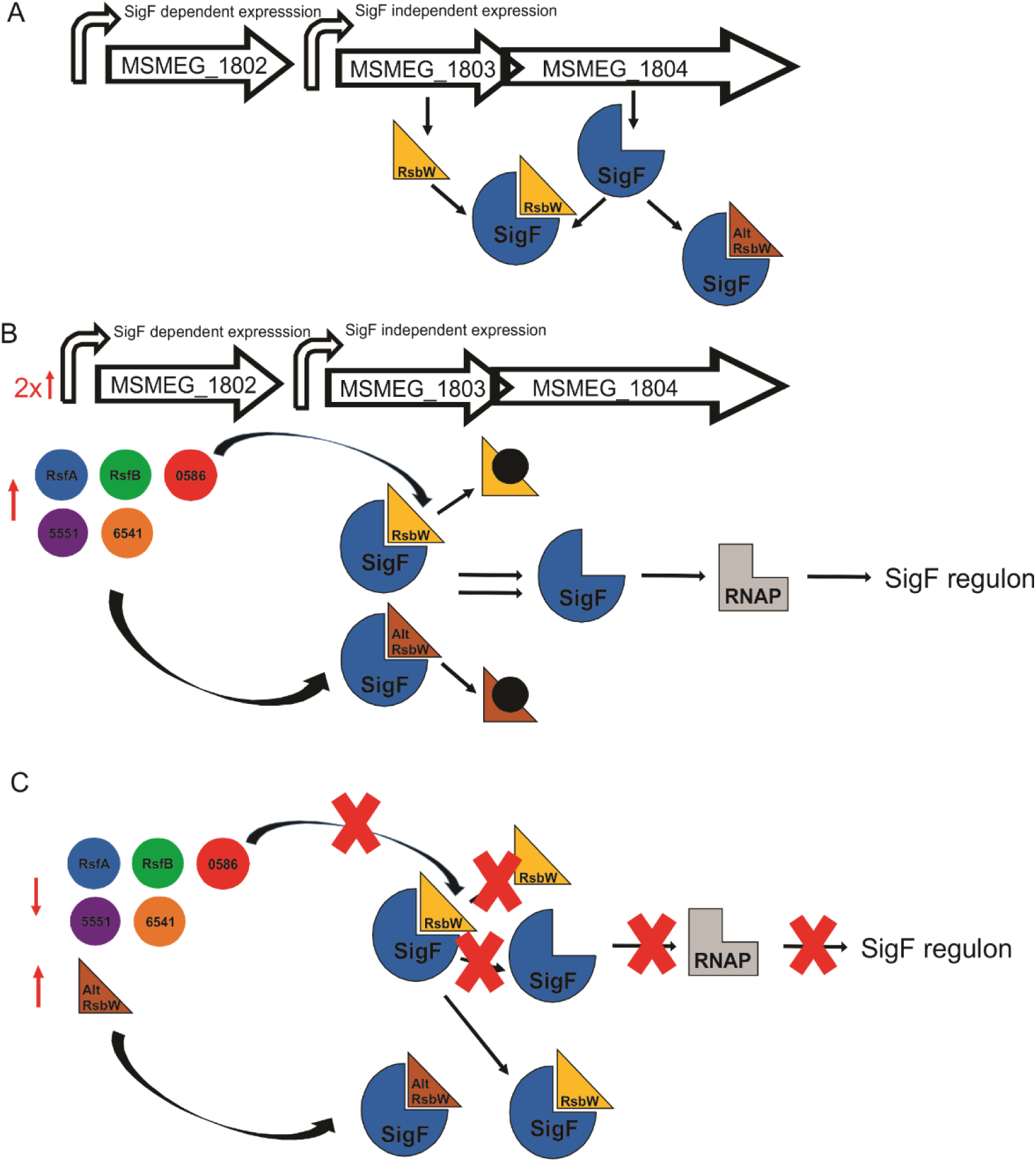
Model of the predicted SigF regulatory cascade. (A) The *sigF* gene in *M. smegmatis* is co-expressed with *rsbW* and *chaB* from two promoter sites. Expression from the promoter located upstream of *chaB* is dependent on SigF whereas expression from the second promoter upstream of *rsbW* is independent and constitutive throughout growth [6]. Under normal growth, SigF activity is sequestered by co-expression of its cognate anti-sigma factor RsbW. The second copy of RsbW (MSMEG_1787) in the *M. smegmatis* genome may further inhibit the activity of SigF. (B) Upon entry into stationary phase the expression of the *sigF* transcript increases 2-fold from the promoter site located upstream of *chaB* [18]. In response to entry into stationary phase and/or to environmental stresses, we hypothesise that expression of one or more anti-sigma factor antagonists increases and that these interact with either RsbW anti-sigma factor protein in a negative fashion, lifting the repression of SigF and permitting its interaction with RNAP. (C) Upon relief of stress, we predict that the expression of anti-sigma factor antagonists decreases, enabling RsbW proteins to remain bound to SigF, sequestering its activity until further stress conditions are met.

The activity of SigF in *M. tuberculosis* is post-translationally regulated by anti-sigma factor UsfX [9] along with two anti-sigma factor antagonists, RsfA and RsfB that further regulate UsfX in a negative fashion [9]. It is likely that the activity of constitutively expressed SigF, transcribed with its cognate anti-sigma factor (UsfX), is sequestered by UsfX, and its repression lifted after release of UsfX upon the expression of the anti-sigma factor antagonists (eg. RsfA and/or B) that are regulated by changing physiological conditions [9]. This permits SigF to bind to RNAP and direct expression of the SigF regulon genes.

In *M. smegmatis*, RsbW (MSMEG_1803) has been shown to interact with SigF [7] suggesting anti-sigma factor function. In addition, a second copy of a *rsbW* gene, (MSMEG_1787), is encoded in the genome and may pose as a second anti-sigma factor of SigF. Further, in *M. smegmatis* (similar to *M. tuberculosis* [19]) there are other predicted anti-sigma factor antagonists [6]. Two, RsfA (MSMEG_1786) and RsfB (MSMEG_6127) have been shown to interact with RsbW [7] and if these function similarly to their counterparts in *M. tuberculosis*, RsfA and B are regulated by two different environmental cues, redox potential and phosphorylation respectively [9]. The remaining three anti-sigma factor antagonists; MSMEG_0586, MSMEG_5551 and MSMEG_6541, comparable to RsfA and RsfB, contain STAS domains (sulfate transporter and anti-sigma factor antagonist domains) and therefore may interact with RsbW.

Collectively, we hypothesise that under normal growth of *M. smegmatis*, constitutively expressed SigF is post-translationally regulated by its co-expressed cognate anti-sigma factor, RsbW, or alternate RsbW (MSMEG_1787) that sequesters the activity of SigF, preventing its interaction with RNAP (Fig 2A). Upon entry into stationary phase, heat or oxidative stress, anti-sigma factor antagonists (RsfA, RsfB, MSMEG_0586, MSMEG_5551, MSMEG_6541) lift the repression caused by either RsbW (MSMEG_1803 or MSMEG_1787) permitting SigF to bind to RNAP and direct transcription of SigF regulon genes (Fig 2B). The suite of anti-sigma factor antagonists likely act to lift repression of SigF under different physiological conditions permitting activation of SigF under different stress conditions. Upon relief of stress, the expression of anti-sigma factor antagonists decreases enabling RsbW to bind to SigF, preventing its activity, until further stress is encountered (Fig 2C). Our proposed model of the SigF regulatory cascade suggests tightknit regulation of SigF and fine-tuning of the regulon.

In *ΔphoH2* we observed an increase in the expression profiles of predominantly SigF regulon genes and those associated with the predicted SigF regulatory cascade (File S2). Fig 3 illustrates the changes in gene expression (up or down) of genes that belong to the predicted SigF regulatory cascade in the absence of *phoH2* at 48 hours. The expression of *sigF* transcript genes and *chaB* increased, and the expression of anti-sigma factor antagonist MSMEG_0586 decreased.

**Fig 3:**
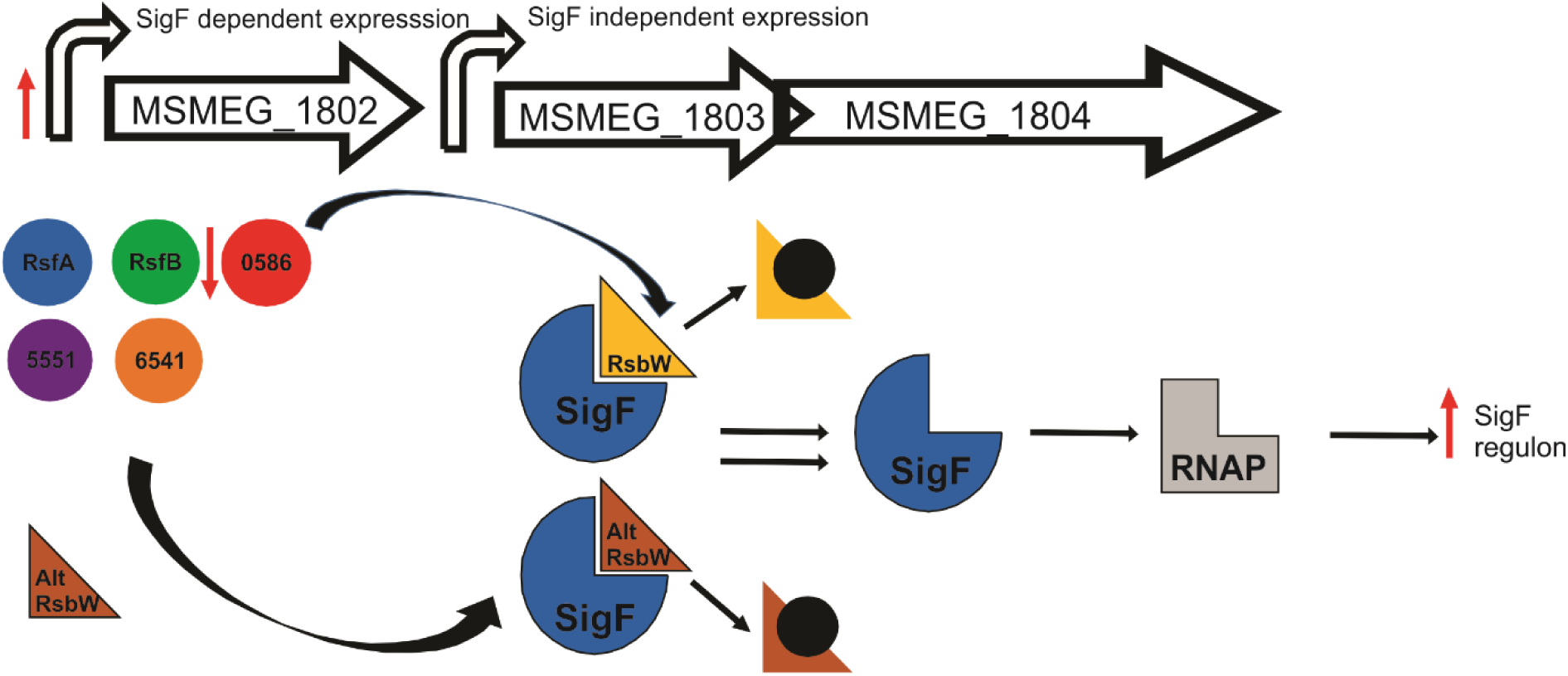
Model of the predicted SigF regulatory cascade and the changes in gene expression observed in *M. smegmatis mc*^*2*^*155 ΔphoH2* compared with *M. smegmatis mc*^*2*^*155*. Red arrows indicate the direction of change in expression (up or down) of genes in *M. smegmatis mc*^*2*^*155 ΔphoH2* in relation to *M. smegmatis mc*^*2*^*155* at 48 hours of growth that met shortlist criteria. The expression of *sigF* transcript genes and SigF regulon genes increased in the absence of PhoH2.

This suggests that PhoH2 contributes to the finely tuned regulation of SigF as observed by dis-regulation of the expression of SigF regulon genes, the *sigF* mRNA transcript and members of the predicted regulatory cascade, under normal growth conditions, in the absence of PhoH2. The remaining genes that comprise the predicted regulatory cascade that did not meet the stringent shortlist criteria, did meet the criteria for DEG (FDR ≤ 0.001 and an absolute value of Log2 ratio of ≥1). Genes here that were upregulated included MSMEG_1803 (RsbW), MSMEG_1787 (RsbW), anti-sigma factor antagonists MSMEG_6541 and MSMEG_5551, and those down regulated included MSMEG_1786 (RsfA) and MSMEG_6127 (RsfB) plausibly suggesting further fine tuning by PhoH2. MSMEG_1804 (SigF) expression was elevated at 48 hours but not greater than Log2 ratio of ≥2.

### PhoH2 targets *sigF* RNA

We hypothesise that PhoH2 targets *sigF* mRNA by way of its targeted mRNAse-helicase activity to fine-tune transcript levels during normal growth. To test this hypothesis, biologically two *sigF* transcripts; *rsbW-sigF* and *sigF* were amplified from *M. smegmatis mc*^*2*^*155* genomic DNA and transcribed into RNA using MEGAscript T7 transcription kit. These were presented as targets, for unwinding and degradation by PhoH2, in reactions over a time-course. RNA transcribed from MSMEG_0467 was used as a negative control to confirm the specificity of PhoH2 for *sigF* transcripts. This gene did not show changes in its expression profile between *M. smegmatis mc*^*2*^*155* and *M. smegmatis mc*^*2*^*155 ΔphoH2* at 48 hours and was of similar length (740 bases) and GC content (64.4 %) to the target RNA sequences. Fig 4 shows the unwinding and degradation of *sigF* RNA (Fig 4A) as shown by the decrease in RNA intensity and increase in smearing of the RNA over time in the presence of PhoH2. Activity was also observed on *rsbW-sigF* RNA (Fig 4B). No activity was seen on MSMEG_0467 RNA (Fig 4C) or by PhoH2-R339A (PhoH-domain mutant with no helicase activity) when used in place of PhoH2.

**Fig 4:**
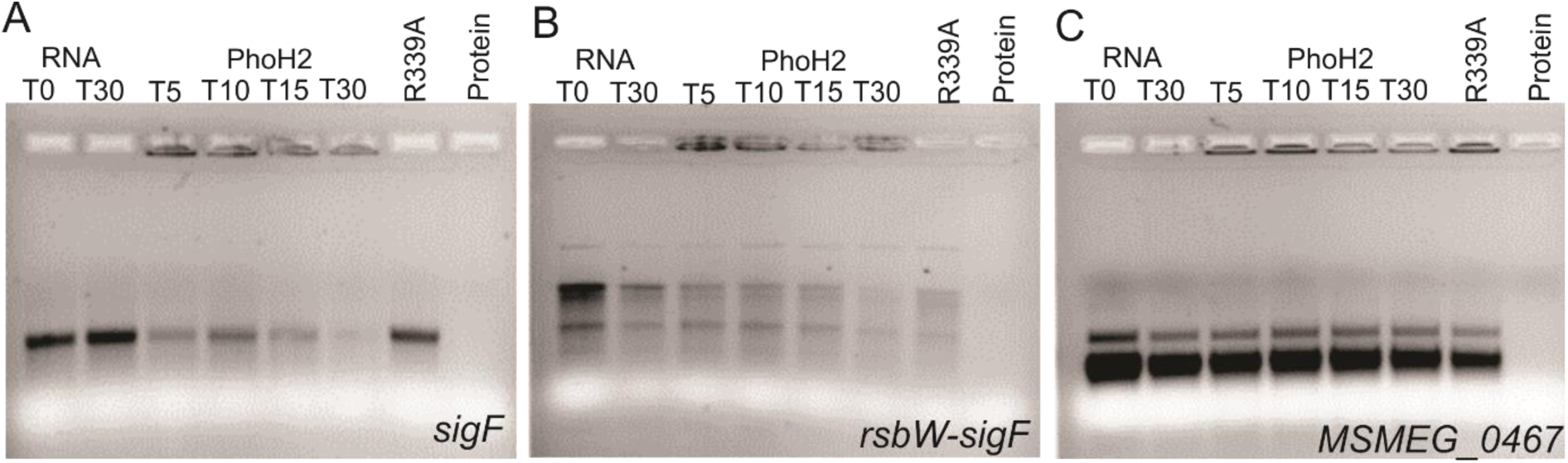
PhoH2 targets *sigF* RNA. (A) *sigF*, (B) *rsbW-sigF* and (C) *MSMEG_0467* RNA transcripts were assayed with PhoH2 over time. RNA unwinding and degradation by PhoH2 of *sigF* RNA is shown by the decrease in RNA intensity and increase in smearing of the RNA over time in the presence of PhoH2. No activity was observed by PhoH2-R339A (RNA helicase mutant) when used in place of PhoH2 nor on MSMEG_0467 RNA C). Reactions/labels: RNA only at T_0_ and T_30_ -RNA (25 µM) incubated in reactions containing 5 mM ATP made up to 15 µl with assay buffer (50 mM tris pH 7.5 20 mM NaCl 5 mM MgCl_2_) at 37 °C. T_5_, T_15_, T_30_ -RNA + PhoH2 (125 µM) - incubated in reactions as above for 5, 15 or 30 minutes. R339A - RNA + PhoH2-R339A in place of PhoH2. Protein - protein only. Reactions were analysed by 1.5 % TAE electrophoresis.

## Discussion

This is the first report experimentally validating the biological target of PhoH2. Three studies report on the possible function of PhoH2 [13-15]. One on PhoH2s involvement in regulating the infection process of *Synechococcus sp. strain WH8102* by Cyanomyovirus [13]. Another on PhoH2s involvement in the response of *C. glutamicum* to limiting phosphate [14] and evidence for a decrease in the expression of *phoH2* during transition from exponential growth to stationary phase growth of *C. glutamicum* [15]. This work suggested that this phase of growth is modulated by SigB [15] and infers that *phoH2* expression is not under the control of SigB.

PhoH2 has sequence-specific RNA and ATP-dependent activity, with preference for double-stranded RNA with 5’-3’ overhangs, and the terminal combination of RNA bases 5’-A C [A/U] [A/U] [G/C] [10]. Each of the RNA transcripts tested here contain versions of this preferred sequence and the greatest number identified in lie in the *sigF* transcript (Table 2 and Fig 5).

**Table 2:** Preferred target sequences and their occurrence in *sigF* transcripts.

| Sequence | Upstream *chaB* | *chaB* | Intergenic | *rsbW* | *sigF* | MSMEG_0467 |
| --- | --- | --- | --- | --- | --- | --- |
| ACAAG |  | 1 | 1 |  |  |  |
| ACAAC |  |  |  |  | 4 | 1 |
| ACAUG | 1 |  |  |  | 3 |  |
| ACAUC |  | 1 |  |  | 1 |  |
| ACUAG |  |  |  |  |  |  |
| ACUUG |  |  |  |  | 1 | 1 |
| ACUUC |  |  |  | 1 | 2 |  |
| ACUAC |  |  |  |  |  |  |
| **Total** | 1 | 2 | 1 | 1 | 11 | 2 |

**Fig 5:**
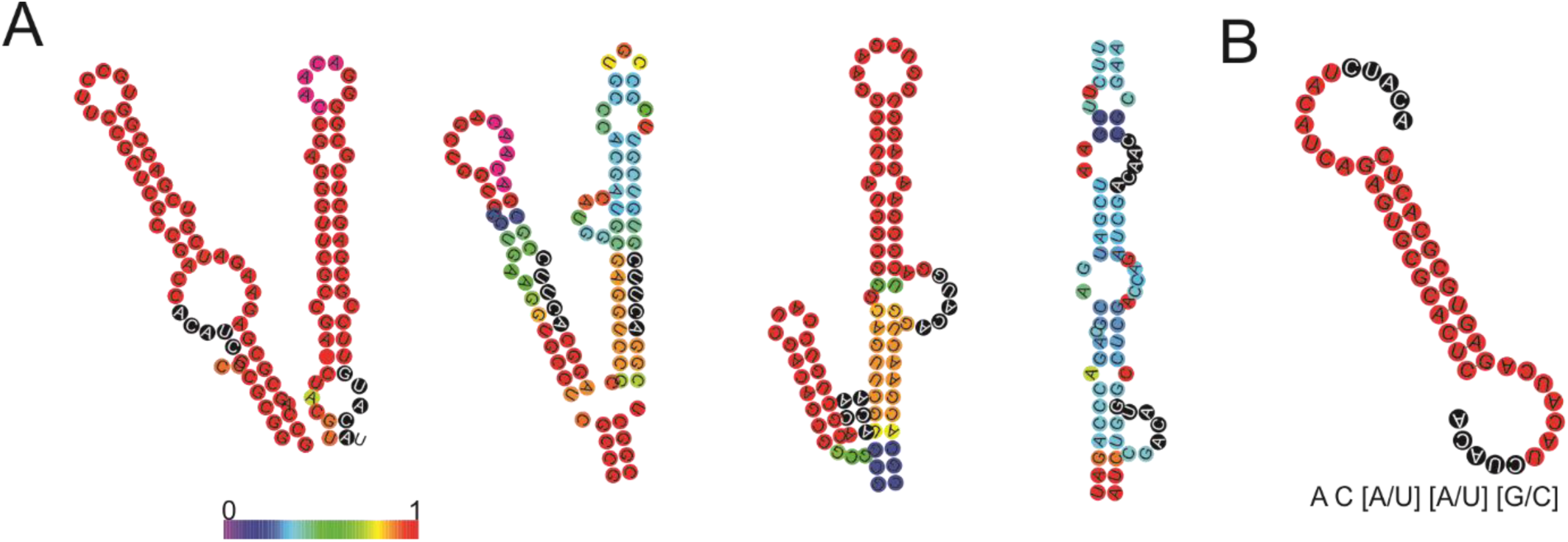
Preferred target sites identified in *sigF* are positioned within stems and loops. (A) Four major stem-loop secondary structures taken from the predicted secondary structure of *sigF* RNA and (B) predicted secondary structure of 5’5 RNA and the preferred target sequence [11]. The stem loops are coloured by base pair probability and the preferred target sequences are highlighted with black circles and white text. RNA secondary structure was modelled by RNAfold [23].

The locations of these target sites are distributed throughout *sigF* mRNA most frequently occurring in loops (9 of the 11 sites) (Fig 5). This observation compliments the *in vitro* biochemical activity of PhoH2 that is specific to single-stranded RNA [10] and suggests that *in vivo*, the PhoH2 hexamer assembles on RNA loops. The mechanism of substrate binding and translocation has been determined for the related hexameric RNA helicase Rho [20]. Rho encircles single-strands of RNA and tethers RNA via its Q and R loops [20]. These loops are responsible for interactions with incoming substrate and project a spiral staircase into the central pore of the hexamer [20, 21]. The Q and R loop staircase recognises and tracks the phosphodiester backbone of RNA, and in conjunction with sequential firing of the asymmetric subunits of the helicase ring, that are in different ATP binding states (nucleotide exchange, ATP-bound, hydrolysis competent and product state), together pull the phosphates and sugars through the ring [20, 21].

Evidence thus far suggests that PhoH2 adopts a hexameric quaternary structure [22]. The active site forming between neighbouring subunits, positioning the nucleic acid binding motifs (RRB1 and RRB2) adjacent to one another on loops near the central pore (Fig 6). The reported inherent flexibility of hexameric helicase subunits [20] may enable these motifs to project into the pore, reminiscent of the Q and R loops of Rho, facilitating RNA recognition and threading into the pore. The unwound RNA may then be fed into and degraded by the PIN-domain RNAse ring. This mechanism implies general RNAse activity of the PIN-domain of PhoH2 and may explain the conservation of PhoAT, the small protein expressed with PhoH2, that may function to sequester RNAse activity, reminiscent of VapB of VapBC toxin-antitoxin systems.

**Fig 6:**
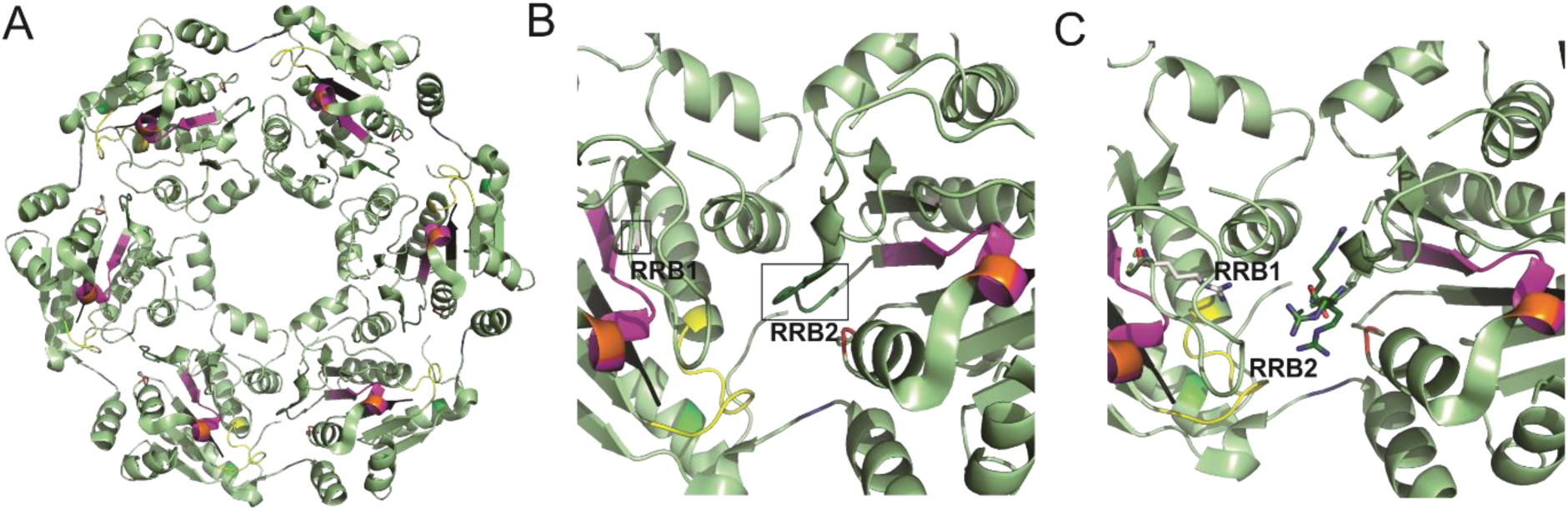
Hexameric structure and active site of the PhoH RNA helicase. Hexameric structure of the PhoH domain of PhoH2 (A) showing the central pore and position of conserved motifs: Q motif (green), Walker A (yellow), Recognition and binding 1 (RRB1) (light grey), RRB2 (dark green), Walker B (magenta), Sensor I (orange), Second Region of Homology (red), Motif III (black) and Sensor II (blue) [10]. (B) Magnified view of the PhoH active site that forms between two adjacent monomers. The positions of RRB1 and 2 are highlighted (boxed) and located on loops. (C) Magnified view of PhoH2 active site showing the position of highly conserved arginine residues (shown as sticks) critical for RNA recognition and binding. Images were made in PyMol V2X using PDB 3B85.

We propose that PhoH2, by way of its targeted RNA unwinding and RNAse activity, contributes, on a post-transcriptional level, to the regulation of SigF by modulating transcript levels. We predict that during growth, constitutively expressed *rsbW-sigF* leads to a pool of RsbW-SigF in the cell. Upon entry into stationary phase there is an increase in transcription from the promoter site located upstream of *chaB* [18] resulting in increased expression of *chaB-rsbW-sigF* mRNA. Depending on the physiological change, we expect increased expression of anti-sigma factor antagonist mRNA in order to lift the repression of RsbW, permitting SigF to bind to RNAP and initiate transcription of its regulon as well as increasing the expression of its own mRNA transcript [18]. Upon established stationary phase and/or alleviation of stress, the expression of anti-sigma factor antagonists is decreased, permitting RsbW to remain bound to SigF, sequestering its activity. We propose that PhoH2 functions to moderate the pool of *sigF* mRNA, and so modulating the SigF response. With controlled levels of *sigF* mRNA and SigF in the cells, this allows for fine control of the cells response to changing physiological conditions. The further upregulation of alternate RsbW (MSMEG_1787) and down regulation of MSMEG_0586 (predicted anti-sigma factor antagonist) in the absence of PhoH2 further supports a role of PhoH2 in SigF regulatory cascade modulation.

These results add further complexity and provide the first report of a post-transcriptional mechanism of regulation of SigF in mycobacteria. This mechanism likely operates in conjunction with the post-translational mechanisms of regulation to enable tightknit control of both SigF and its regulon of genes through the transition from exponential to stationary phase of growth and under changing environmental conditions.

## Acknowledgements

The Waikato Medical Research Foundation and University of Waikato Strategic Initiative Fund for consumables funding.

## Supporting information

**S1 Fig: PCR confirmation of *phoH2* deletion from *M. smegmatis mc***^***2***^***155.***

**S1 Table: Primers used in this study.**

**S1 File. Biological target RNA sequences.**

**S2 File. Summary of sequencing and alignment statistics and list of differentially expressed genes (DEGS).**

**S1 Fig:**
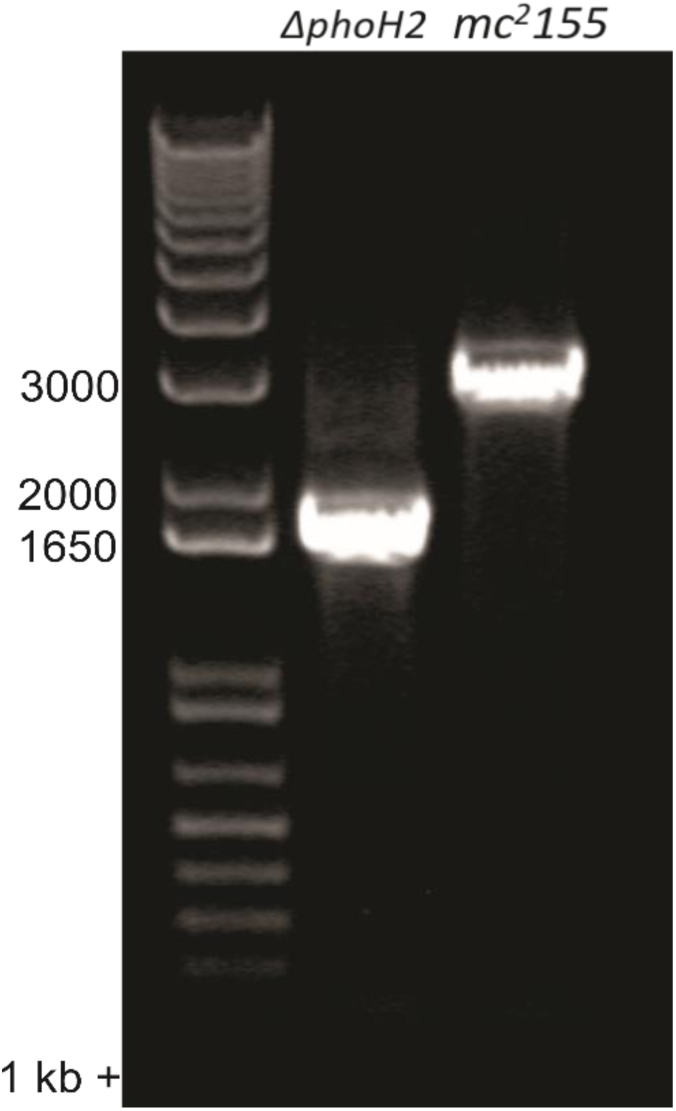
PCR confirmation of *phoH2* deletion from *M. smegmatis mc*^*2*^*155.* PCR using primers that flanked the deletion site were used with *M. smegmatis mc*^*2*^*155* or mc^2^155 *ΔphoH2* DNA as template to confirm the deletion of *phoH2*. Expected band sizes: *mc*^*2*^*155* – 2914 bp and *mc*^*2*^*155 ΔphoH2* – 1622 bp.

**Supplementary Table S1:**
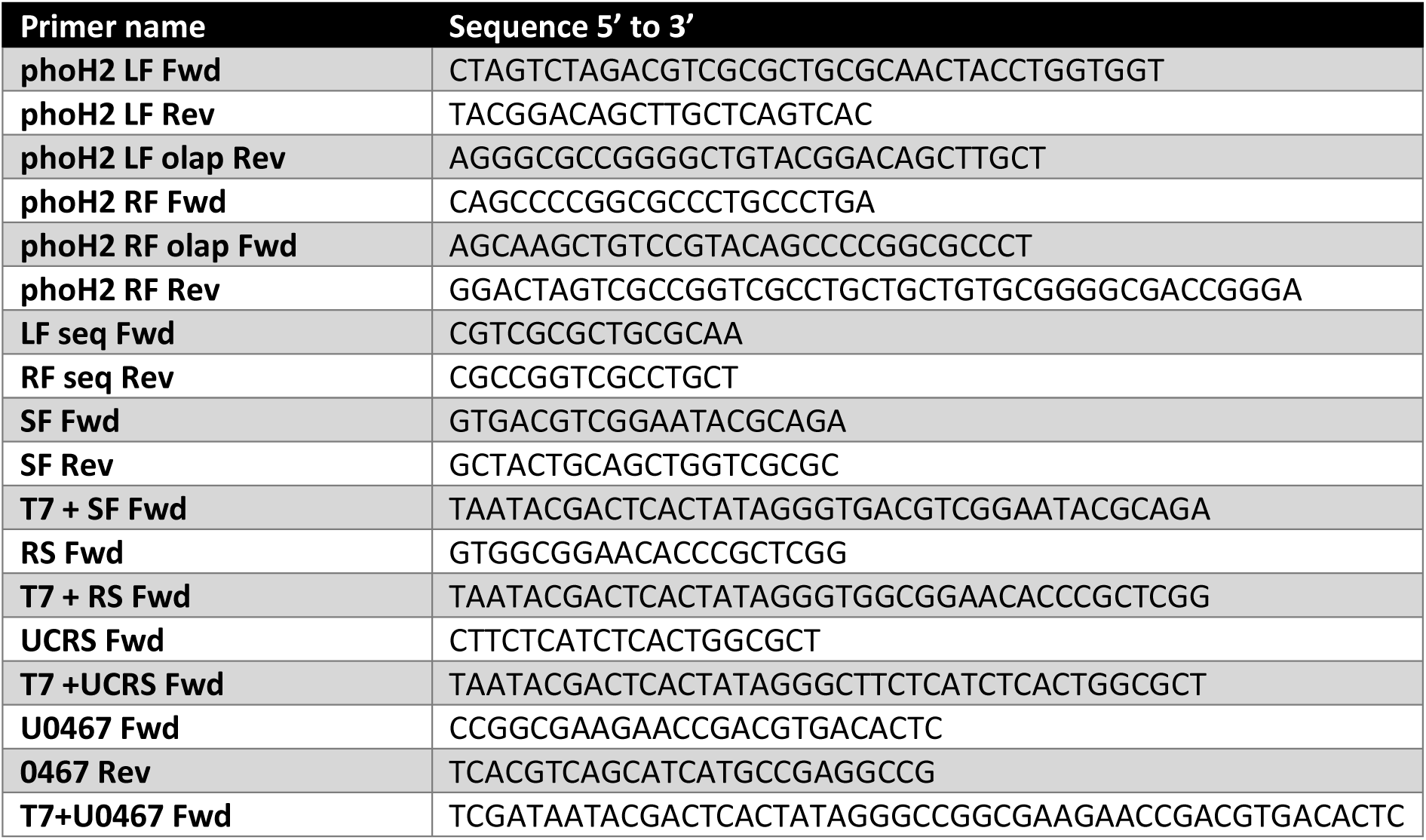
Primers used in this study.

**File S1. Biological target RNA sequences**

**Bold** - Translational start site

<u>Underlined</u> - preferred target

*Upstream + chaB + intergenic + rsbW + sigF* (1889 bp)

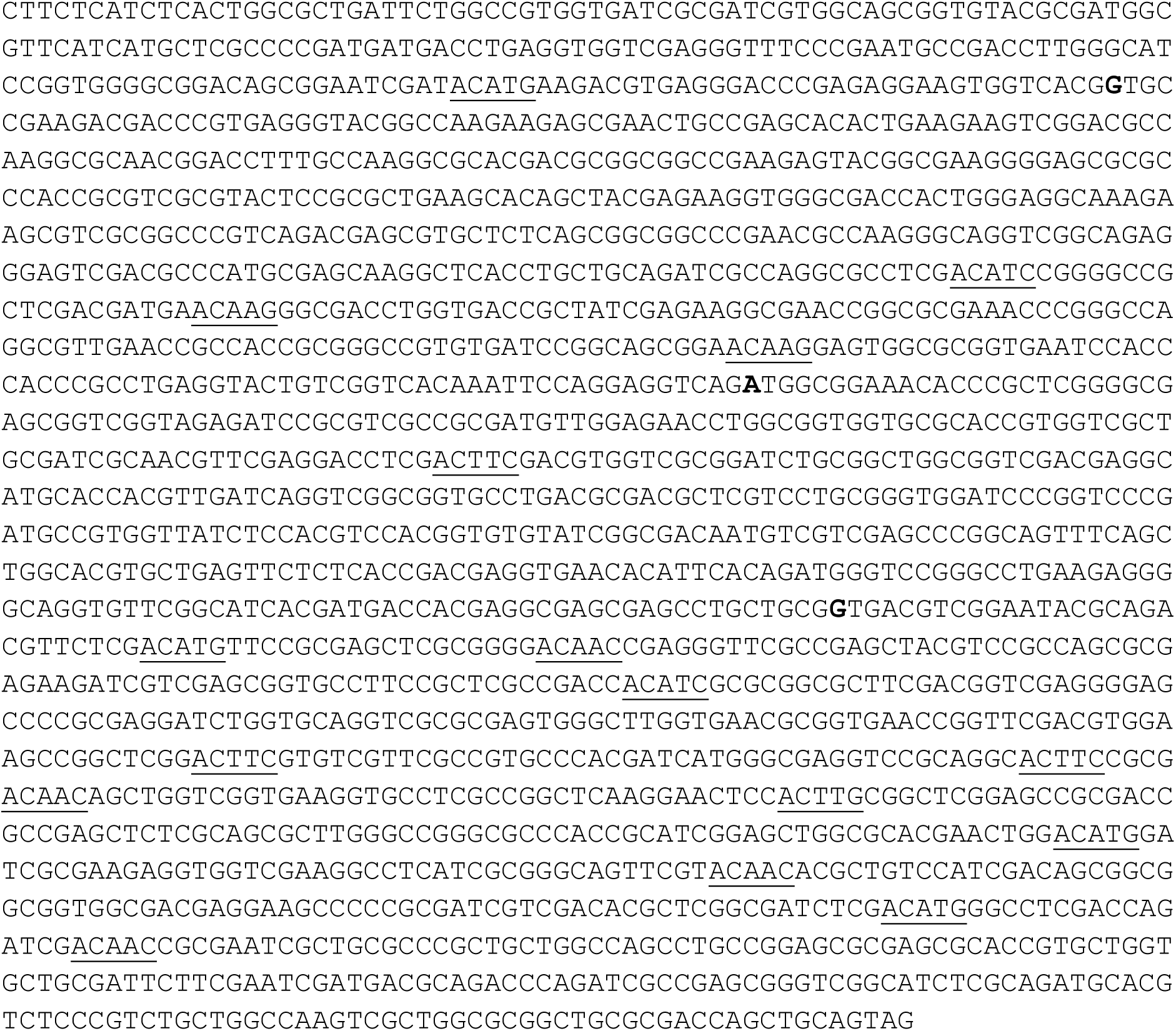

*rsbW + sigF* (1166 bp)

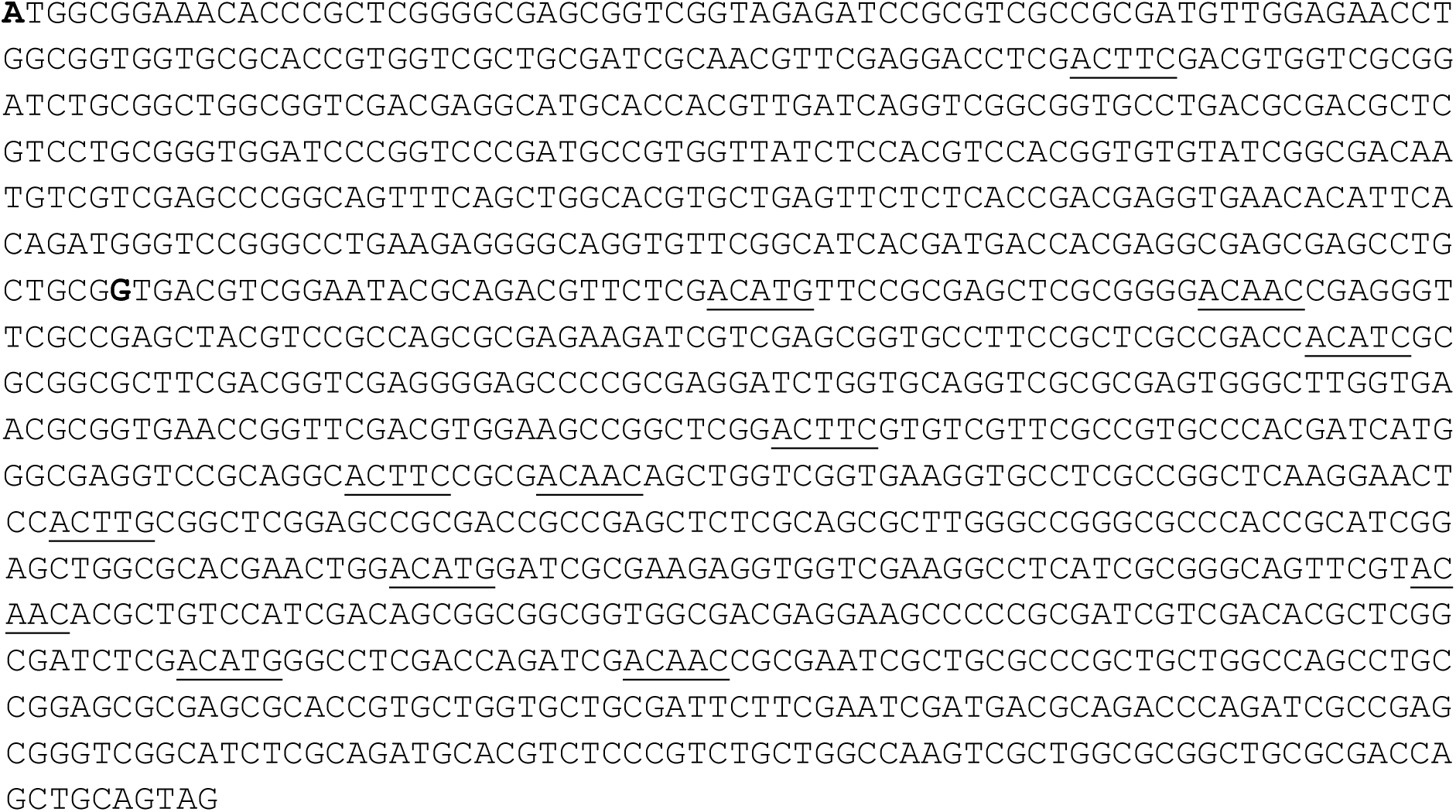

*sigF* (753 bp)

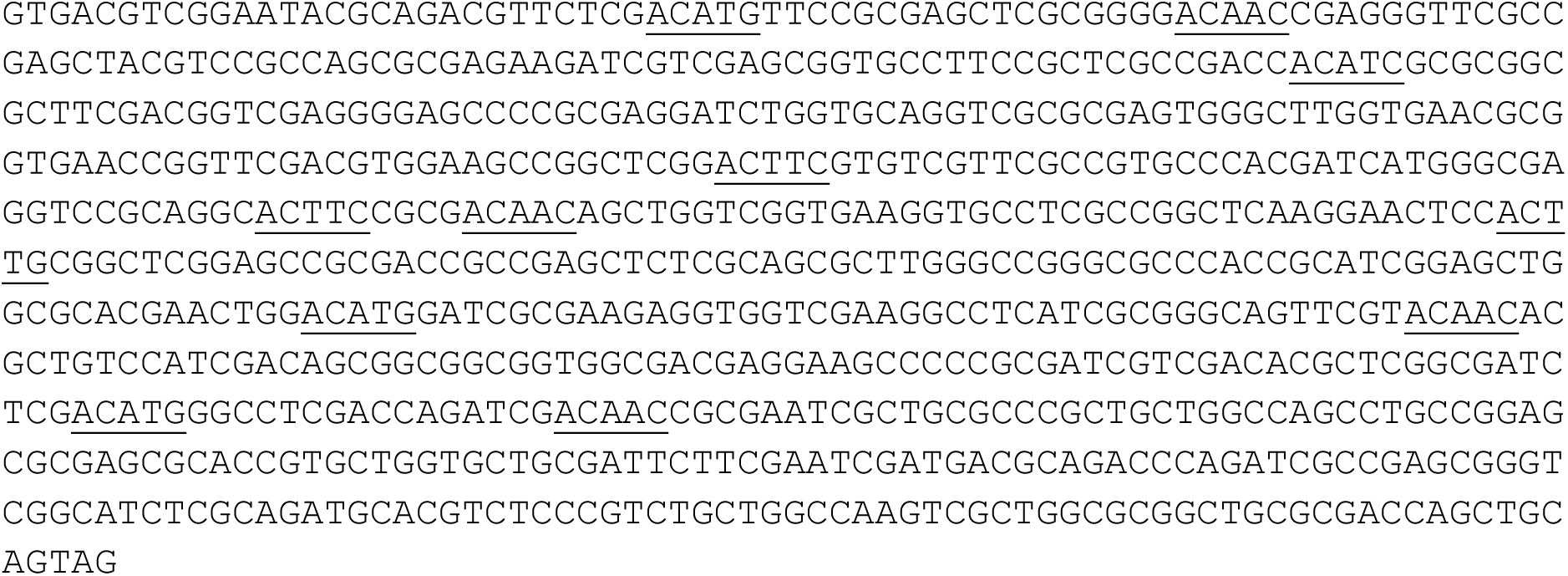

*Negative control MSMEG_0467 (with upstream)* (740 bp)

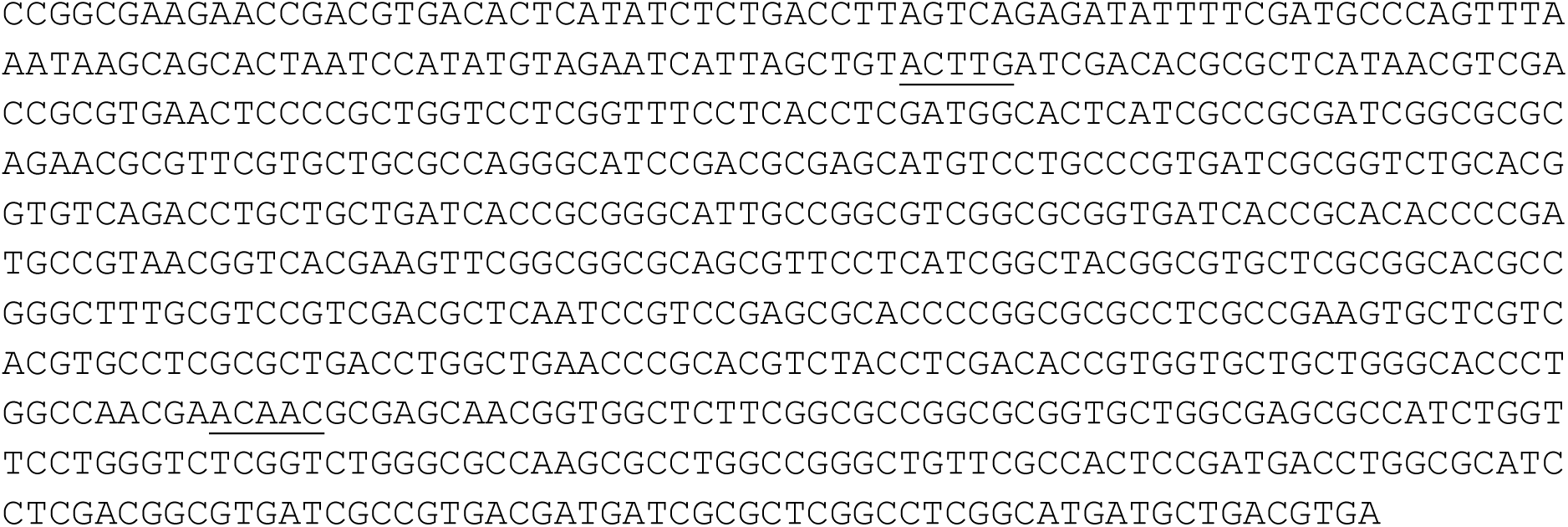

**File S2:**
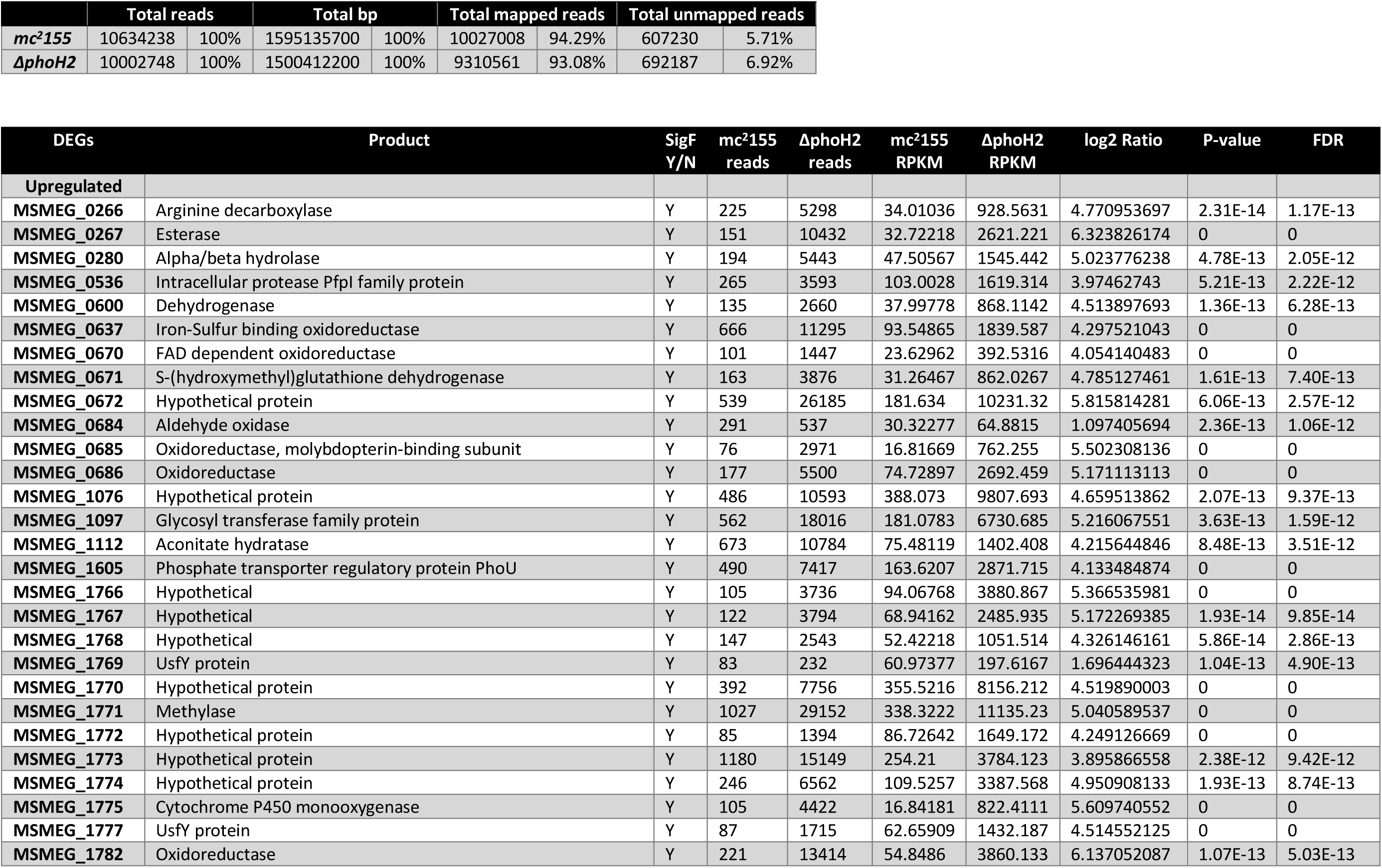
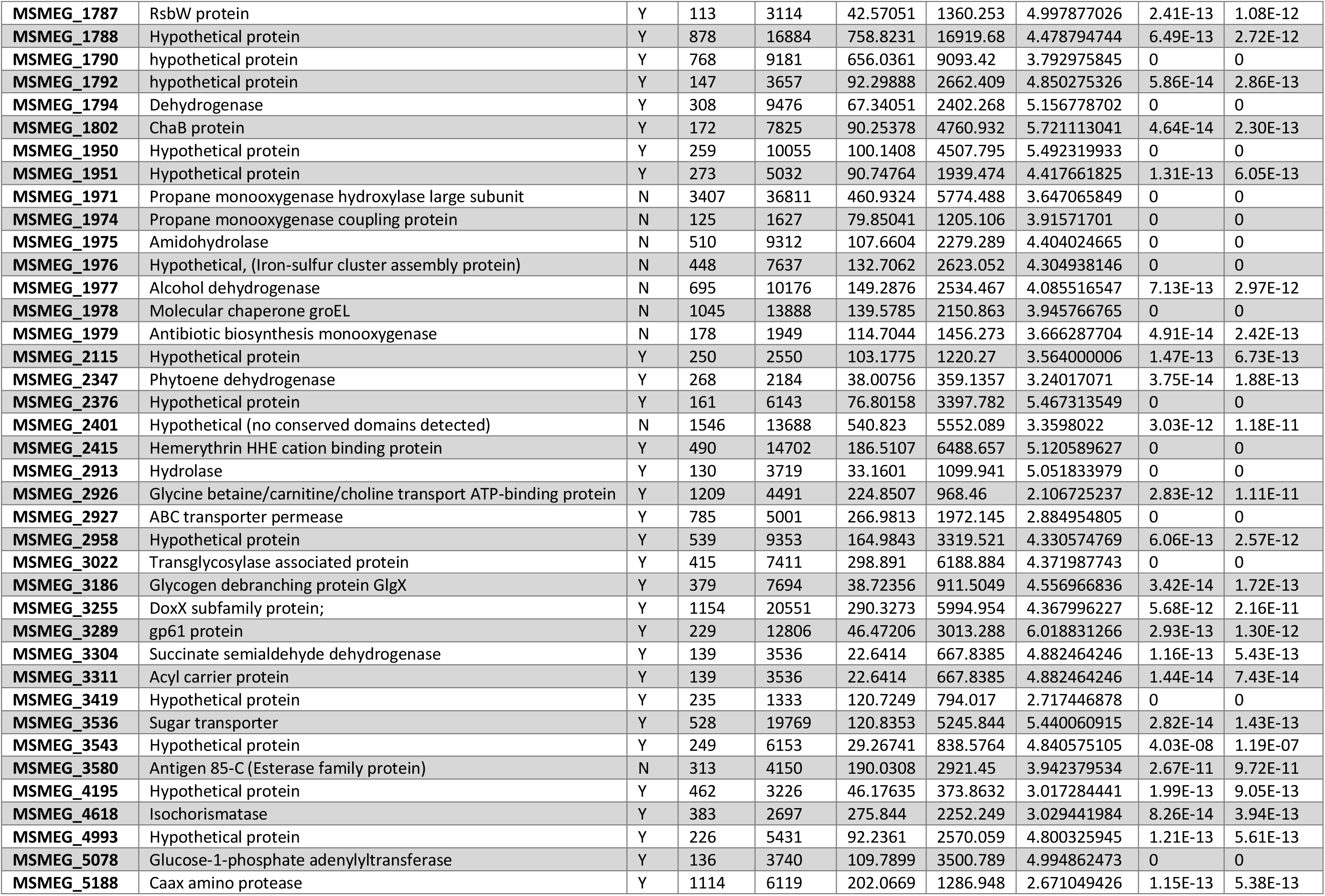
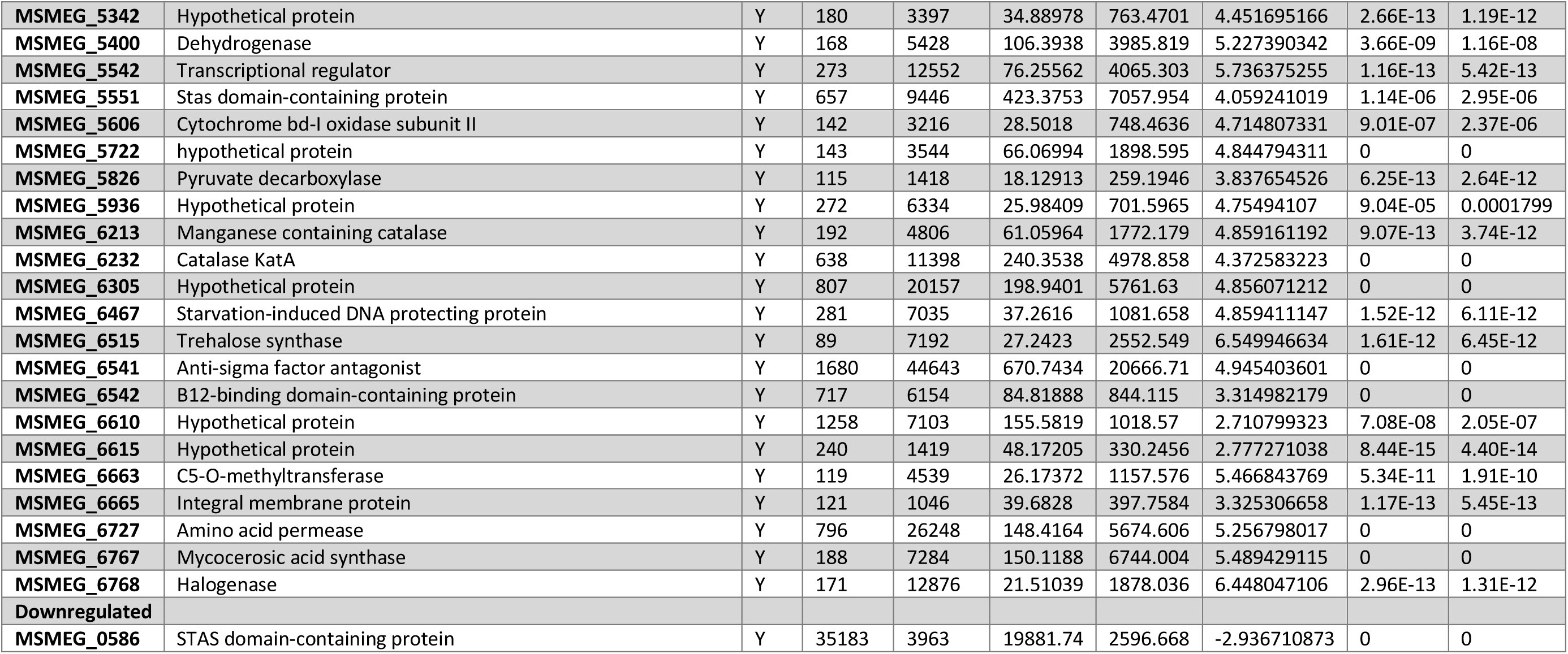
Summary of sequencing statistics and list of differentially expressed genes (DEGS) (FDR ≤ 0.001 ≥2 Log2 ratio ≥75 reads)

